# Age-dependence and aging-dependence: The case of neuronal loss and lifespan in a *C. elegans* model of Parkinson’s disease

**DOI:** 10.1101/098038

**Authors:** Javier Apfeld, Walter Fontana

**Affiliations:** Biology Department, Northeastern University, Boston MA 02115, USA; Department of Systems Biology, Harvard Medical School, Boston MA 02115, USA

## Abstract

It is often assumed, but not established, that the major neurodegenerative diseases, such as Parkinson’s disease, are not just age-dependent (their incidence changes with time) but actually aging-dependent (their incidence is coupled to the process that determines lifespan). To determine a dependence on the aging process requires the joint probability distribution of disease onset and lifespan. For human Parkinson’s disease, such a joint distribution is not available because the disease cuts lifespan short. To acquire a joint distribution, we resorted to an established *C. elegans* model of Parkinson’s disease in which the loss of dopaminergic neurons is not fatal. We find that lifespan is not correlated with the loss of neurons and that a lifespan-extending intervention into insulin/IGF1 signaling accelerates neuronal loss, while leaving death and neuronal loss times uncorrelated. This suggests that distinct and compartmentalized instances of the same genetically encoded insulin/IGF1 signaling machinery act independently to control neurodegeneration and lifespan in *C. elegans*. Although the human context might well be different, our study calls attention to maintaining a rigorous distinction between age-dependence and aging-dependence.

## INTRODUCTION

In demographic parlance, the term “age” refers to the time elapsed since the occurrence of a specific developmental event, for example the onset of adulthood. To conform with this meaning of age, the phrase “age-dependent disease” should simply refer to a disease whose risk of onset (or progression to any particular stage) changes with time, usually in a steadily increasing fashion as is the case for a variety of metabolic and neurodegenerative diseases as well as many cancers^1^. Yet, the phrase is often used in a way that appears to carry the connotation of age as a “biological time” marking the process of aging, effectively suggesting an *aging*-dependence and not just an age-dependence. Such slippage amounts to asserting a causal connection between aging and (say) the onset of a disease. The significance of such a connection derives from the hope that interventions that slow aging might delay disease onset. The main point of our contribution is to show that this frequent slippage from age-dependence to aging-dependence is all but warranted.

Aging refers to a process that (typically) increases the risk of death with the passing of time, i.e. age. The risk of death in turn determines the lifespan distribution and the survival function. This means that the risk of death depends not only on time—which makes it age-dependent—but outright defines aging—which makes it trivially aging-dependent. Having defined aging, it is clear that to determine whether disease X or more precisely any phase of disease X, such as its onset, depends on aging, one has to study the co-variation of disease X and lifespan. This involves first determining the joint probability distribution of the time of onset of X and the time of death *from aging*, and then assessing, whether or not this joint probability distribution is the product of the respective marginal distributions. A difficulty arises when the progression of an age-dependent *and lethal* disease caps lifespan. This is known in statistics as a ceiling effect. If an individual dies from a disease, we cannot know how long that individual would have lived had the disease not killed but allowed the aging process to take its course. This counterfactual prevents us from acquiring a joint probability distribution and seems to have impeded a rigorous determination of whether any age-dependent disease with fatal outcome is actually dependent on the aging process. In the United States, such diseases include the seven leading causes of death among individuals aged 65 years and over, accounting for more than 70% of deaths in this age group^1^. The wide-spread hope of delaying the onset of such diseases by intervening in the aging process betrays a tacit assumption that a dependence on aging must exist.

One way to probe the connection between age- and aging-dependence is to focus on diseases that are not lethal, which might be less pressing. Another approach, and the one we chose, is to exploit the transfer of a disease that is lethal in humans to a model organism in which it is not. Here we aim at a basic analysis of the age- and aging-dependence of a Parkinson’s disease model in the nematode *C. elegans.* While this does not permit inference to the human case, it nonetheless illustrates the interaction between aging and a model of a time-dependent physiological condition in an organismic context that has a history of illuminating molecular processes in humans.

Parkinson’s disease is perhaps one of the most widely studied age-dependent diseases. In humans, the incidence of Parkinson’s disease increases with age at a nearly exponential rate, with a mean age at onset of 70 years^2^. The pathogenesis of Parkinson’s disease is characterized by the loss of dopaminergic neurons in the substantia nigra of the brain^3^. The progression of Parkinson’s disease involves the loss of essential neurons, leading to the early death of affected individuals^4^. A well-validated *C*. *elegans* Parkinson’s disease model has been used previously to identify several genetic determinants of dopaminergic neurodegeneration^5,6^, including a gene responsible for familial forms of Parkinson’s disease in humans^7^. However, unlike in humans, dopaminergic neurons are not essential for survival in *C*. *elegans*^8^. The disease model provided us therefore with a unique platform to study the poorly understood connections between age-dependent neurodegeneration and the aging process that determines the organism’s lifespan.

The *C*. *elegans* model is significant also because its lifespan is controlled by widely-conserved mechanisms. Early studies in *C*. *elegans* showed that lifespan can be more than doubled by reduction-of-function mutations in the insulin/IGF1 receptor gene *daf*-*2*^9^ or in its downstream signaling effectors, including the *age*-*1* phosphatidylinositol 3-kinase (PI3K) gene^10^. Subsequent studies showed that lowering the action of insulin and IGF1 receptors also extends lifespan in the fruit fly *Drosophila melanogaster*^11,12^ and in mice^13,14^. In worms and flies these lifespan-extending effects are mediated by the activation of FOXO transcription factors, DAF-16^15,16^ and dFOXO^17,18^, respectively. This functional conservation appears to extend to humans, since *Foxo3A* gene variants are associated with human longevity^19,20^.

Reducing insulin/IGF1 receptor signaling or increasing FOXO activity specifically in adipose or neuroendocrine tissue can increase lifespan in worms^21–23^, flies^18,24^ and mice^13,25^. This indicates that, across phylogeny, insulin/IGF1 signaling regulates lifespan, at least in part, through the action of systemic signals generated by these tissues. These signals are thought to regulate the cellular processes responsible for the functional decline associated with aging and the associated increase in the risk of death. As such, they may represent ideal entry points for the development of therapies against aging-dependent diseases^26^.

We began our study by investigating the role of insulin/IGF1 signaling in the regulation of the timing of the loss of dopaminergic neurons in the *C*. *elegans* Parkinson’s disease model. We found, unexpectedly, that long-lived insulin/IGF1 pathway mutants exhibit a more severe degenerative phenotype. These effects are mediated by the action of *daf*-*16*; but we also found that this transcription factor regulates neurodegeneration *independently* from lifespan, suggesting constraints to the organism-wide action of the insulin/IGF1 pathway. We then probed the joint probability distribution of the timing of neuronal loss and the death of the organism in the Parkinson’s disease model. We found that the onset of neurodegeneration and aging (as manifest in the timing of death) are independent phenomena, yet influenced by the action of distinct and independent insulin/IGF1-dependent processes coexisting in a compartmentalized fashion within the same organism.

## RESULTS

### Insulin/IGF1 signaling affects age-dependent dopaminergic degeneration

In order to investigate the role of insulin/IGF1 signaling in the control of the timing of neuronal degeneration, we used a well-validated *C*. *elegans* transgenic model of Parkinson’s disease^5,6^. In this model, human α-synuclein and green fluorescent protein (GFP) are co-expressed in the eight dopaminergic neurons of *C*. *elegans* hermaphrodites^5,6^. GFP enables *in vivo* visualization of these neurons by fluorescence microscopy^27^. α-synuclein is a protein that plays a central role in the etiology and progression of Parkinson’s disease^28^. An increase in the α-synuclein gene dosage causes early-onset forms of familial Parkinson’s disease in humans^29^. Similarly, expression of human α-synuclein in dopaminergic neurons of *C*. *elegans* causes age-dependent neurodegeneration^5,6^. This degeneration is mediated by the worm homologue of the familial Parkinson’s disease gene *PARK9/ATP13A2*, suggesting that the mechanisms leading to the pathological effects of α-synuclein are evolutionarily conserved^7^.

We characterized the age-dependent degeneration of four dopaminergic neurons located in the head of the animal, the so-called cephalic neurons (two dorsal and two ventral) in the *C. elegans* Parkinson’s disease model^5,6^ by examining the neurodegenerative phenotype in larvae and adults of increasing age. The oldest animals we examined, day-7 adults, had lived two thirds of their average expected lifespan. They were post-reproductive and tended to exhibit the reduced and uncoordinated locomotion characteristic of old age^30^. We found that larvae and day-0 adults never exhibit neurodegeneration in any of the four cephalic neurons (Fig. 1 and Table S1). However, as animals grew older, their cephalic neurons were more likely to be absent (Fig. 1). Each of the cephalic neurons exhibits this age-dependent neurodegenerative phenotype, although it is more severe in the ventral pair than in the dorsal pair. The neuronal-loss phenotype could be the result of neuronal death or of neuronal dysfunction leading to GFP expression levels below the detection limit. In any case, this phenotype requires α-synuclein expression, as we did not observe a loss of cephalic neurons at any age in animals that express only GFP in these neurons (Table S1).

**Figure 1:**
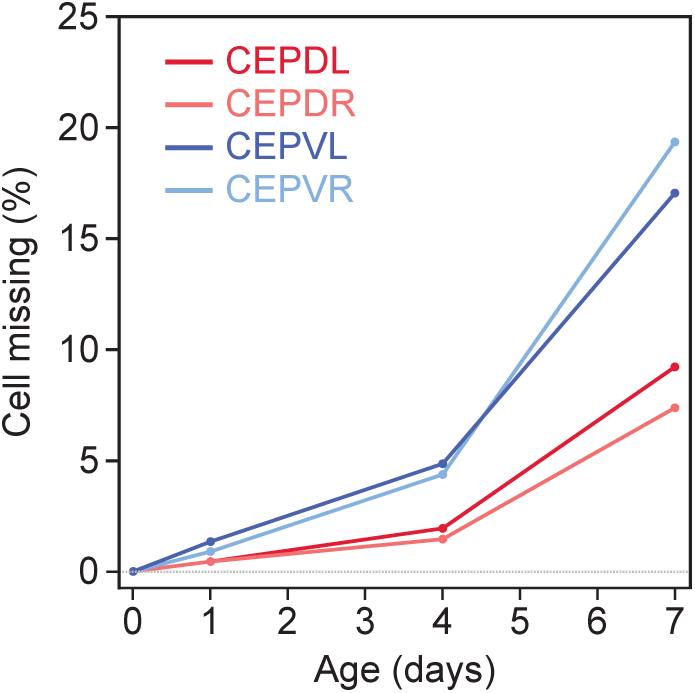
Age-dependent loss of cephalic neurons in the *C*. *elegans* Parkinson’s disease model. Age-dependent loss of cephalic neurons in animals expressing α-synuclein in the dopaminergic neurons (Parkinson’s disease model). *C*. *elegans* has four cephalic (CEP) neurons, located dorsally (D) or ventrally (V), on the left (L) or the right (R) side of the worm. Ordinal logistic regression indicates that the proportion of animals with an absent neuron increases in an age-dependent manner (*P* = 0.0009 between days 1 and 4, and *P* < 0.0001 between days 4 and 7) and that dorsal neurons have a more severe phenotype than ventral neurons (*P* < 0.0001), while left-right position has no impact on phenotype (*P* > 0.05). Animals that do not express α-synuclein do not exhibit neuronal loss (Table S1). For additional statistics see Table S1.

To determine whether insulin signaling affects the timing of neuronal loss in the Parkinson’s disease model, we tested the effect of the *age*-*1(hx546)* reduction-of-function mutation. As expected, in the Parkinson’s disease model this mutation causes a large, 2.6 fold, lifespan extension (Fig. 2B and Table S2). Surprisingly, these long-lived mutants exhibit a more severe neurodegenerative phenotype than wild-type animals: the fraction of missing ventral cephalic neurons in seven-day old *age*-*1(hx546)* mutants is much higher than in wild-type animals (Fig. 2A and Table S3). In contrast, the fraction of missing dorsal cephalic neurons is not affected by *age*-*1(hx546)* (Fig. 2A and Table S3). In *daf*-*2(e1370)* mutants, where insulin/IGF1 signaling is more strongly reduced than in *age*-*1(hx546)* mutants^31,32^, we also observed increased loss of ventral cephalic neurons in seven-day old animals (Table S4). In addition, the fraction of missing dorsal cephalic neurons in seven-day old *daf*-*2(e1370)* mutants is lower than in wild-type animals (Table S4). We conclude that insulin/IGF1 signaling, a modulator of aging, has opposite and cell-type specific effects on the age-dependent loss of dopaminergic neurons in the *C*. *elegans* Parkinson’s disease model.

**Figure 2:**
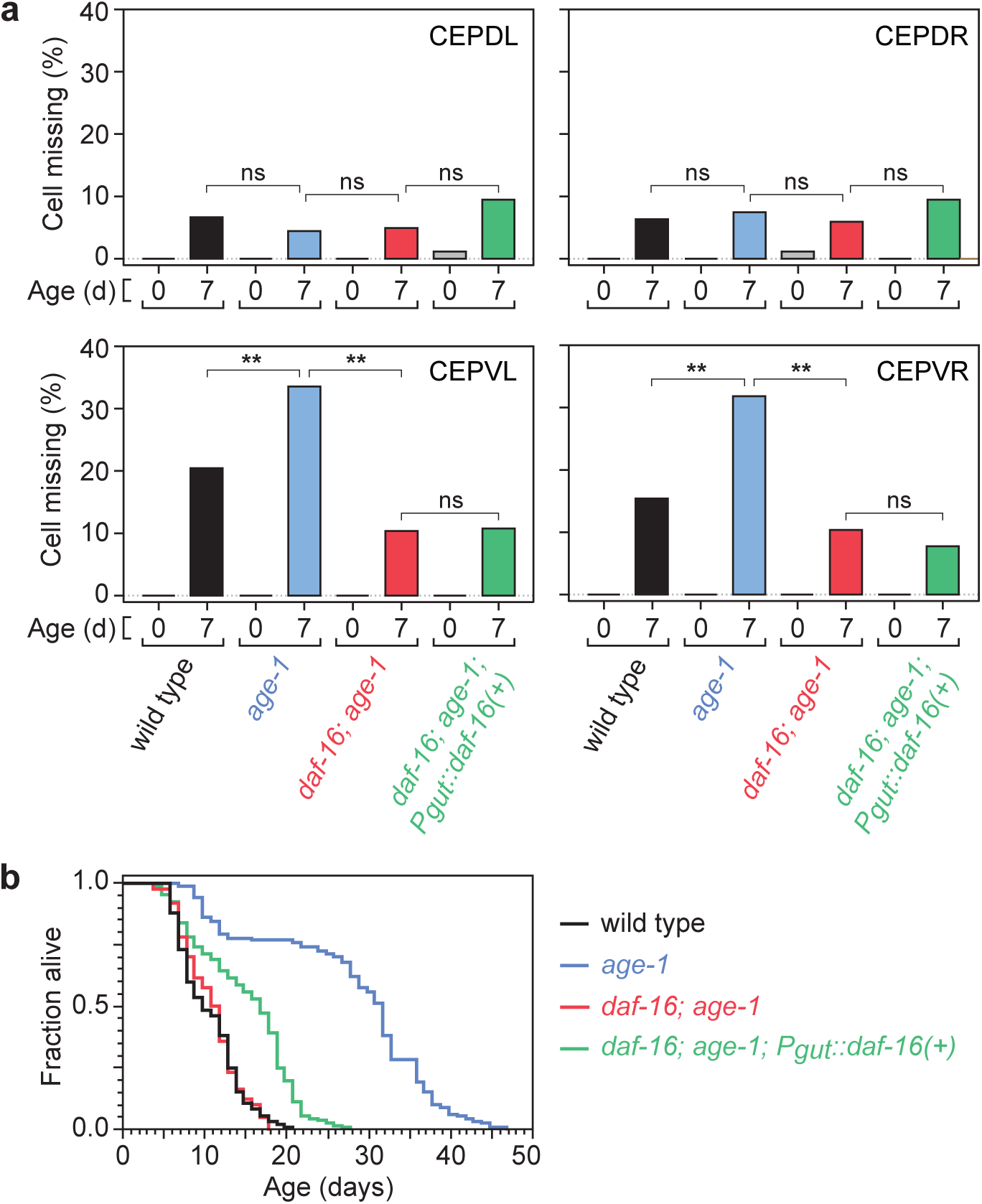
Effect of insulin/IGF1 signaling on neurodegeneration and lifespan. (**A**) Each panel shows the percentage of animals with the indicated cephalic neuron missing in young adults (day 0) and old adults (day 7). These percentages are paired for each specified genotype. Except for *age*-*1* (CEPDL) and *daf*-*16*; *age*-*1* (CEPDL and CEPDR), all pairs show a statistically significant difference of at least *P* = 0.004 between young and old. The double asterisk (**) indicates *P* ≤ 0.001; “ns” indicates that no significant difference was observed (*P* > 0.05). For additional statistics see Table S1. (**B**) Survival curves for wild-type and mutant animals expressing α-synuclein in the dopaminergic neurons (Parkinson’s disease model). For statistics see Table S2.

Activation of FOXO transcription factors mediates the lifespan-extending effects of reduced insulin/IGF-1 signaling in *C*. *elegans* and *Drosophila*^9,33–35.^ We therefore asked whether the sole *C. elegans* FOXO transcription factor^15,16^, DAF-16, plays a similar role in mediating the effects of reduced insulin/IGF1 signaling on neuronal loss by examining the effect of the *daf*-*16(mu86)* null allele^15^. The observed fraction of missing ventral cephalic neurons was dramatically lower in seven-day old *daf*-*16(mu86)*; *age*-*1(hx546)* double mutants than in *age*-*1(hx546)* single mutants (Fig. 2A and Table S3). Therefore, *daf*-*16* is required for the increased loss of ventral cephalic neurons observed in *age*-*1(hx546)* mutants. Our findings, thus, indicate that activation of *daf*-*16* mediates both lifespan extending and neurodegenerative phenotypes associated with reduced insulin/IGF1 signaling. We conclude that the divergent effects of this pathway on the timing of neurodegeneration and lifespan are generated downstream of the action of *daf*-*16*.

### Distinct tissues affect neurodegeneration and lifespan through insulin/IGF1 signaling

Tissue-restricted activation of FOXO transcription factors is sufficient to extend lifespan in *C*. *elegans*^22,36^ and *Drosophila*^17,18,24,37.^ In *C. elegans,* FOXO activation in one tissue can influence gene expression in other tissues via cell-to-cell signaling mechanisms that involve *daf*-*16* in the receiving tissue (FOXO-to-FOXO signaling) and by signaling mechanisms that do not (FOXO-to-other signaling)^22,23,36,38.^ Currently it is not known whether FOXO-to-FOXO signaling is an important determinant of nematode lifespan, but FOXO-to-other signaling does play a role in the modulation of lifespan, since *daf*-*16* expression in the intestine is sufficient to extend lifespan in animals that lack *daf-16* in all other tissues^22,23^. Importantly, FOXO-to-other signaling from the intestine also delays the age-dependent paralysis caused by muscle-specific expression of the human Alzheimer’s protein Aβ_1-42_^36^. We therefore asked whether this type of systemic signaling mechanism also affects the timing of neuronal loss in the Parkinson’s disease model.

We found that expressing *daf*-*16(*+) only in the intestine extends the lifespans of *daf*-*1*6*(mu86)*; *age*-*1(hx546)* double mutants by 40% in the Parkinson’s disease model (Fig. 2B and Table S2). However, this treatment does not affect the loss of any of the four cephalic neurons on seven-day old adults (Fig. 2A and Table S3). Hence, tissues other than the intestine must mediate the effect of insulin/IGF1 signaling on α-synuclein induced neuronal loss.

### Lifespan and neurodegeneration are the outcomes of independent processes

We have shown that mutations affecting insulin/IGF1 signaling components are able to influence both the timing of neuronal loss and lifespan. We therefore decided to determine whether a coupling between the timing of neuronal loss and of death occurs in the disease model. If the timing of α-synuclein induced neuronal loss and of organismic death were indeed coordinated, then neuronal loss would depend on aging, not just age. Specifically, animals in which a particular cephalic neuron is present at a given age would have a different lifespan from those in which that neuron is absent. In addition, the magnitude of the difference in lifespan between these groups would provide information about the sign and strength of the correlation between the (model) disease process and the aging process. We therefore compared the lifespans of animals that experienced neuronal loss with the lifespans of those that did not.

We imaged 3,397 seven-day old adults and determined whether each of the four cephalic neurons was present or absent in each animal. We then recovered each animal and determined its lifespan. We found that animals in which a specific cephalic neuron is present have lifespans that are indistinguishable from those where that neuron is absent (Fig. 3A, Fig. 4A-B, Fig. S1A-D and Table S5). We also do not observe a difference in lifespan among these animals when we group them (i) by whether any of the four cephalic neurons is missing or not, (ii) by the number of missing dorsal cephalic neurons, (iii) by the number of missing ventral cephalic neurons, or (iv) by the total number of missing cephalic neurons (Fig. S2 and Table S5). These findings strongly suggest that the timing of neuronal loss and organismic death are independent.

**Figure 3:**
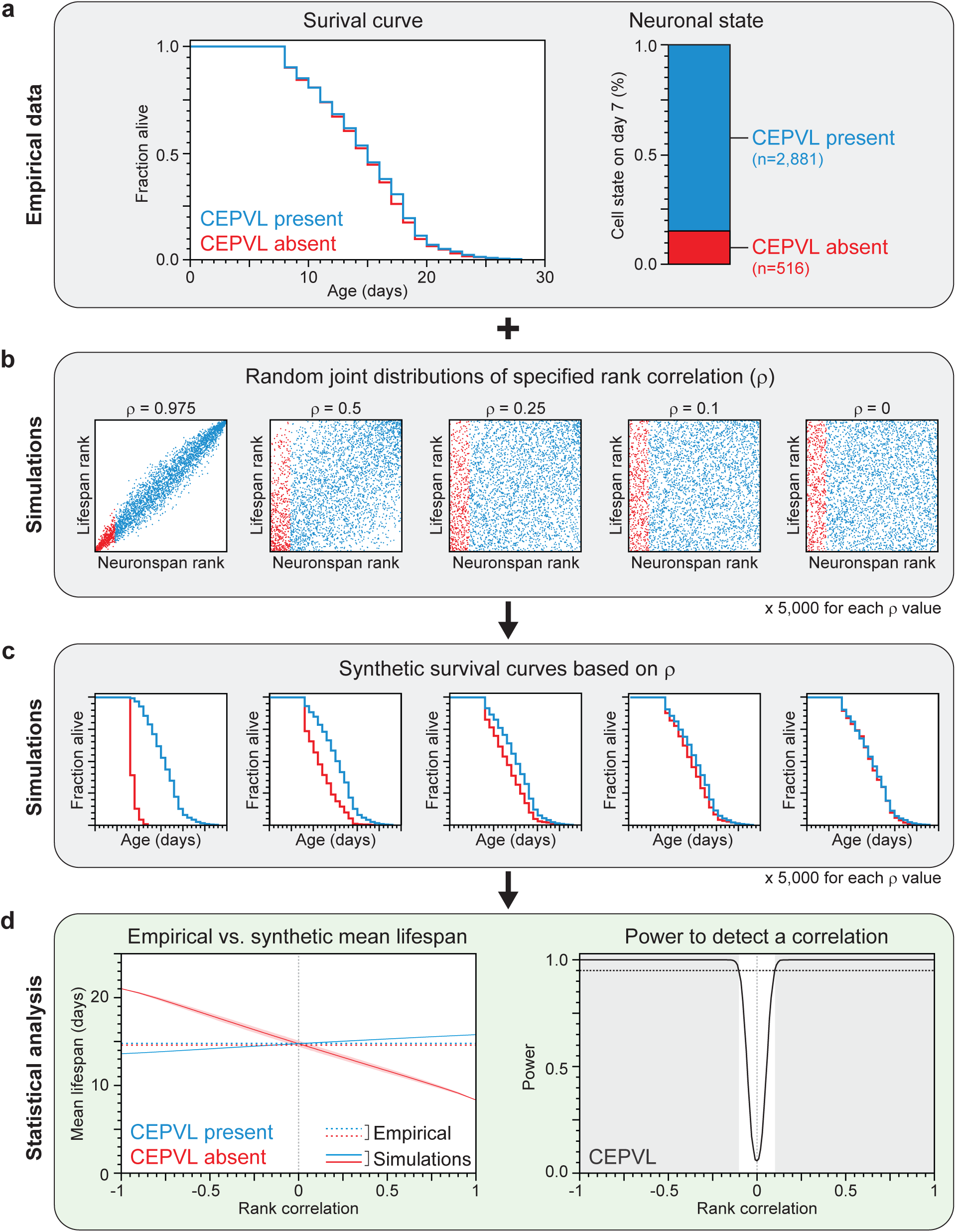
Statistical analysis strategy based on empirical data and computational simulations. (**A**) This panel shows the two types of data that we collected for each individual expressing α-synuclein in the dopaminergic neurons: its lifespan and the state of each of its four cephalic neurons on day 7 of adulthood. These data were used to construct the empirical distribution of lifespan (plotted as a survival curve) and neuronal state. (**B**) We generated collections of random joint distributions of varying Spearman rank correlation values. Each point represents a simulated animal that has a relative lifespan and neuronspan. Simulated animals where a neuron is absent on day 7 have the lowest neuronspans (red points), while those were a neuron was present have the highest neuronspans (blue points). The joint distribution size equals the number of individuals in the empirical data set and the proportion of points labelled as “present” or “absent” equals the observed proportion of neuronal states. (**C**) We generated synthetic survival curves by dividing the empirical lifespans based on their rank, using the simulated lifespan ranks of the present (blue) and absent (red) groups in each random joint distribution. (**D**) The collection of synthetic survival curves was analysed statistically to model how the mean lifespan of each group varies with Spearman rank correlation value. The shaded areas represent the 95% point-wise confidence interval of the mean lifespans of the synthetic curves. The statistical power of our experiment is the fraction of simulations with a statistically significant (*P* < 0.05) difference in survival between the neuron present and absent groups. The shaded region denotes the Spearman’s rank correlation values where statistical power is greater than 95% (dotted line). The empirical data and statistical analysis shown in this figure is also shown in Fig 4B,F,J.

**Figure 4:**
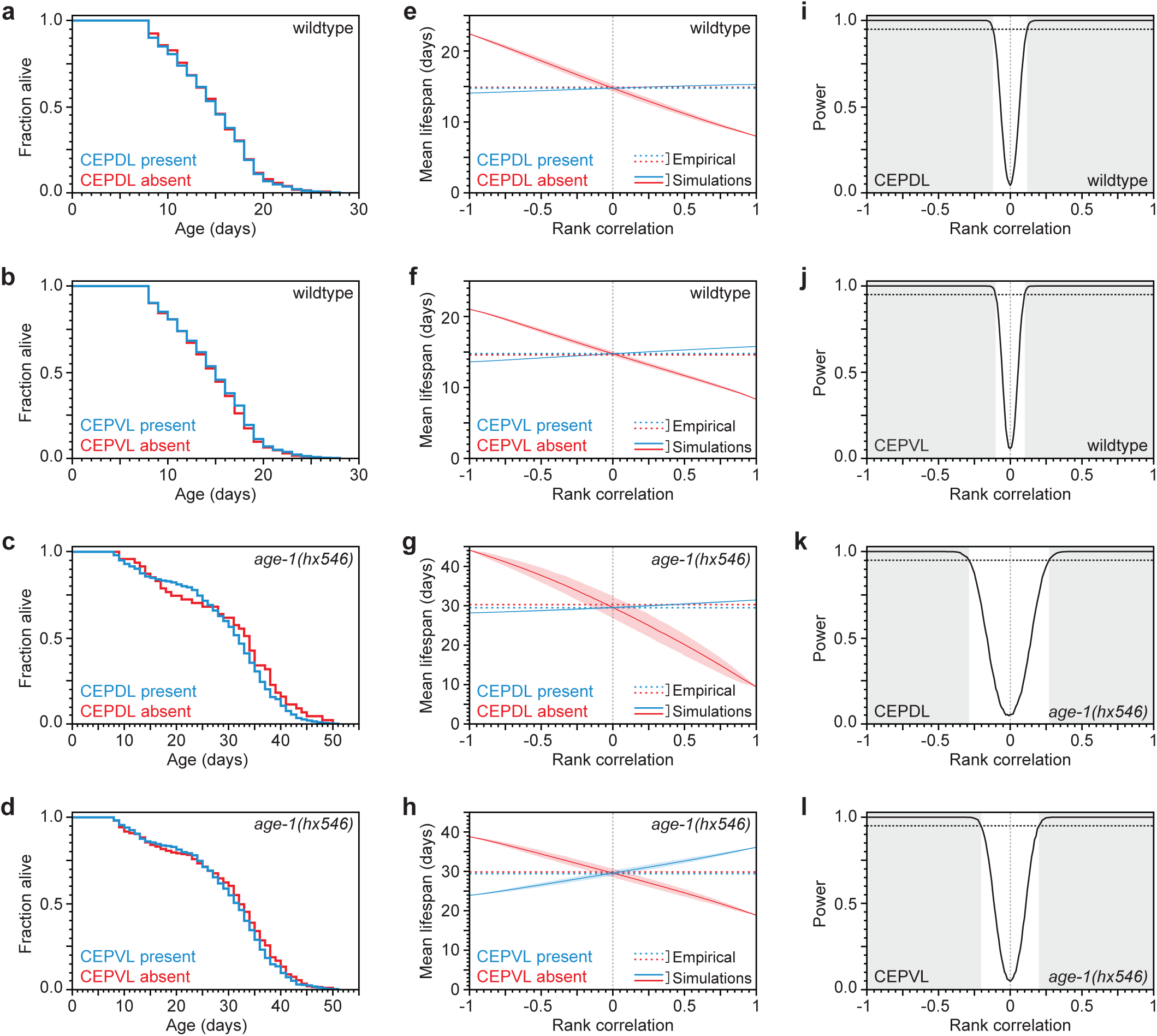
Independence of neuronal loss and organismic death in wild-type and *age*-*1* mutant animals. (**A-D**) No difference in lifespan was observed when 3,397 seven-day old wild-type animals (A-B) or 542 seven-day old *age*-*1(hx546)* mutants (C-D) expressing α-synuclein in the dopaminergic neurons were grouped according to whether the specified cephalic neuron was present (blue line) or absent (red line). For statistics see Table S5. (**E-H**) Using computer simulations (Fig. 3), we predicted how the mean lifespans of seven-day old animals with present (red line) and absent (blue line) cephalic neurons are influenced by the Spearman’s rank correlation between the timing of neuronal loss and lifespan. The shaded areas represent the 95% point-wise confidence interval of the predicted mean lifespans. The observed mean lifespans (dotted lines) are only consistent with correlations of low magnitude. (**I-L**) The statistical power of our experiment is the fraction of simulations with a significant (*P* < 0.05) difference in survival between the neuron present and absent groups. The shaded region denotes the Spearman’s rank correlation values where statistical power is greater than 95% (dotted line). For the analysis of based on the state of CEPDR and CEPVR on day 7 in wild type and *age*-*1* mutant animals see Figs. S3 and S4, respectively.

To ascertain that lifespan distributions (survival functions) in the presence or absence of cephalic neurons are indistinguishable, we tested (using the Wilcoxon test) whether the two survival functions arise from the same underlying distribution, which gave us a *P*-value that the null hypothesis (no statistical difference) holds. Since this result is critical for our thesis that death and neuronal-loss times are uncorrelated, we needed to evaluate whether our experiment had enough power, *i.e.* we wanted to know how often our experiment would yield indistinguishable (*P* > 0.05) survival functions by chance, if there actually was a correlation between death and neuronal-loss times. To assess the statistical power of our experiment, we ideally would like to computationally repeatedly sample 3,397 (the population size in our experiment) death times and neuronal-loss times that are consistent with our empirical data but with varying correlation, then test for statistical equality, and determine, for each correlation value, the frequency with which testing would not miss that there was in fact a correlation. However, in our dataset we do not have a full joint distribution of death times and neuronal-loss times; we only have a death-time distribution and we know the number of individuals that experienced neuronal loss by age (day) 7 (Fig. 3A). We therefore use rank correlations and (for each sample) generate a joint rank distribution of 3,397 points with specified rank correlation (Fig. 3B). We then read off the lifespan ranks for points corresponding up to day 7 of neuronal loss and the lifespan ranks for points from day 7 onward, and convert those ranks back into death times by matching them to the ranks of our empirical lifespan data. This process yields a survival curve each for the subpopulation that has and that has not experienced neuronal loss up to day 7 (Fig. 3C and Movie 1). Finally, we test the two survival curves against a null hypothesis of no statistical difference. In this way, sampling 5,000 joint rank distributions per rank correlation value, we estimate that our experiment is powered to detect a correlation between death and neuronal-loss times greater than 0.13 with probability 0.95 for each of the cephalic neurons (Fig. 3D, Fig. 4E-F,I-J and Fig. S1E-M). Thus, knowing when a specific neuron is lost appears to provide very little, if any, information about organismic lifespan. We conclude that lifespan and neuronal loss are not coordinated by systemic signals; instead, they are the outcomes of independent processes.

Having established independence in the disease model with a wild-type background, we wondered whether reduction of insulin/IGF1 signaling might affect this independence. To test this possibility, we stratified a population of 542 seven-day old *age*-*1(hx546)* adults according to the presence or absence of each cephalic neuron, and then determined their lifespans, following the same scheme as in the wild-type case.

We found that *age*-*1(hx546)* mutants in which a cephalic neuron is present have lifespans that are indistinguishable from those where that neuron is absent (Fig. 4C-D, Fig. S3A-D and Table S5). In addition, we found that the lifespan of *age*-*1(hx546)* mutant individuals is not predicted by the extent of neurodegeneration they exhibited (Fig. S4). As in the wild-type case we carried out numerical simulations to estimate how sampling noise would impact the conclusions we draw from our experiment as a function of rank correlation (Fig. 4G-H,K-L and Fig. S3E-M). This analysis indicates that our experiment was well-powered to detect a significant difference in lifespan for at least one of the four neurons if rank correlations were greater than 0.15 in magnitude (Fig. S3M). We conclude that lifespan and neurodegeneration are at most weakly coordinated, if at all, in *age*-*1(hx546)* mutants.

## DISCUSSION

In this study we probed the causal links between neurodegeneration and lifespan in a *C*. *elegans* model of Parkinson’s disease. We found that while the loss of dopaminergic neurons increases with age, it is phenomenon independent of aging. Yet, despite this independence, the timescales of neuronal loss and organismic lifespan are both regulated by insulin/IGF-1 signaling, pointing to a compartmentalization of this pathway’s action. Our findings have implications for our understanding of a broad class of age-dependent and lethal human diseases whose aging-dependence remains poorly understood.

We first review our observations on the *C*. *elegans* Parkinson’s disease model. We found that specific sets of dopaminergic neurons are affected differently by age. Dorsal cephalic neurons are less likely to exhibit α-synuclein induced loss than ventral cephalic neurons as time progresses. In addition, these neurons are affected differently by insulin/IGF1 signaling. In particular, compared to wild type, *age*-*1/PI3K* mutants exhibit higher neurodegeneration in ventral cephalic neurons but not in dorsal cephalic neurons. In humans, too, though for reasons unknown, specific dopaminergic neurons exhibit distinct susceptibility to Parkinson’s disease^39^. Our findings in *C*. *elegans* suggest that this specificity could stem from differences in the sensitivity of specific dopaminergic neurons to insulin or IGF1.

In addition to extending lifespan in *C*. *elegans*, reducing insulin/IGF1 signaling also delays the onset of a wide variety of phenotypes associated with normal aging, including many biochemical^40,41,^ morphological^30,42,^ and behavioural impairments^9,43,44.^ Neurons are affected too, as in the case of the reduction of the age-dependent incidence of neurite sprouting in ALM and PLM mechanosensory neurons^45-47^. Diminishing the action of the insulin/IGF1 signaling pathway also postpones age-dependent changes in some disease models, including models of age-dependent paralysis due to the expression in body wall muscles of an aggregation-prone polyglutamine-containing protein or the human protein Aβ_1-42_ linked to Alzheimer’s disease^48,49.^ We were therefore surprised to find that reducing insulin/IGF1 signaling augments the loss of ventral cephalic neurons in the Parkinson’s disease model. Previous studies have shown that FOXO-to-other signaling from the intestine affects both lifespan^22^ and the timing of body paralysis due to expression in body wall muscles of the human Aβ_1-42_ protein^36^. In contrast, we find that FOXO-to-other signaling from the intestine does not affect the loss of neurons in the *C*. *elegans* Parkinson’s disease model. Most strikingly, we did not find a correlation between lifespan and neurodegeneration in the disease model with wild-type background.

It is important to emphasize, once again, that age simply refers to physical time whose origin (time zero) is defined by the occurrence of a specific developmental event, which in *C*. *elegans* is the onset of adulthood. However, in informal usage, age often appears to carry a connotation of “biological time”. At the present state of knowledge, any notion of biological time in the context of aging must refer to the risk of death. Thus, the risk of any event X, other than death, is dependent on “biological time” only in so far as it carries information about the risk of death (and vice versa). The risk of death tracks the aging process and the risk of loss of dopaminergic neurons tracks the disease process (or a pathological condition for the worm). Both risks change with time, that is, the age of an individual. An interesting question is whether the disease is not only age-dependent, but also dependent on aging—the very process that determines lifespan. In this study, we showed that the Parkinson’s disease model in *C*. *elegans* is *independent* of aging.

We also found something that might seem puzzling at first: that both processes underlying the respective event risks are influenced by manipulation of insulin/IGF1 signaling (*e.g.* in *age*-*1(hx546)* mutants) and yet remain independent. At the level of abstraction of genetics, the *age*-*1* gene might be viewed as a common cause (all else equal) of its effects on neurodegeneration and aging in the disease model. However, in the context of a fully developed organism, tissue-specific realizations of the signaling pathway may well have distinct effects on distinct processes, in particular if these realizations are “compartmentalized” in the sense of not exchanging state information.

Are age-dependent human diseases aging-dependent? In humans, the progression of Parkinson’s disease and other neurodegenerative diseases often includes the loss of neurons involved in essential functions, thereby resulting in early death^4^. It is therefore not possible to measure the joint probability distribution of Parkinson’s disease onset and lifespan as determined by aging. This has made it difficult to study whether systemic signals coordinate the onset of such diseases with the timing of death from aging. One approach to address this impasse is to use an animal model in which the disease is not fatal, therefore allowing the acquisition of joint probability distributions. This is the case in the *C. elegans* Parkinson’s disease model because, unlike in humans, dopaminergic neurons are not essential for survival in the worm^8^. While the disease model in *C. elegans* might serve well the study of molecular mechanisms that are associated with the disease and potentially conserved in humans, it may fail in evidencing a connection with aging that might be present in humans, perhaps because of a more complex aging process. This said, our results regarding the distinction of age- and aging-dependence suggest at least a cautionary stance with regard to the widely held, yet untested, view that the onset of neurodegenerative diseases, such as Parkinson’s and Alzheimer’s, is mechanistically linked to the aging process that determines lifespan. This assumption is the basis for the expectation that disease onset will be postponed by pharmaceutical, dietary or genetic interventions that extend lifespan^50-53^. Our results call for prudence when using the term “disease of aging”, which is often tacitly based on the notion of an overall aging process that couples all age-related phenotypes—a notion that our work demonstrates does not apply to *C. elegans*.

## METHODS

### General methods and strains

The wild-type strain of *C. elegans* was Bristol N2. Animals were cultured on NGM agar petri plates containing a lawn of *E. coli* OP50 bacteria. Unless otherwise noted, animals were reared at 25°C throughout their lifespan, and were derived from animals reared at 20°C that were transferred to 25°C as young adults (defined as the first 4 hours of adulthood). The following genes and mutations were used: LG I: *daf*-*16(mu86)*; LG II: *age*_-_*1(hx546)*, *fer*-*15(b26)*; LG III: *daf*-*2(e1370)*; LG IV: *baInl1[P_dat_*_-_*_1_::α*-*synuclein*, *P*_dat_*_-_*_1_⸬gfp]*; other: *egIs1*[*P_dat_*_-_*_1_⸬gfp]*; extra-chromosomal: *muEx268[P_ges_*_-_*_1_*⸬gfp*-*daf**-*16(*+), *P*_odr_*_-_*_1_⸬rfp]* (referred to as *P_gut_⸬daf*-*16(*+) in the text)^22^.

### Construction of strains

In double mutant construction, *baInl1* and *egIs1* were assayed by GFP expression in dopaminergic neurons. *daf*-*2(e1370)* and *age*-*1(hx546)* were assayed by the presence of dauers at 25°C and 27°C, respectively. *fer*-*15(b26)* was assayed by the presence of all female progeny at 25°C. *muEx268* was assayed by the presence of RFP expressing neurons. *daf*-*16(mu86)* and *daf*-*16(*+) were distinguished by PCR using allele-specific primers^54^.

### Microscopy

Age-synchronous cohorts of adults were prepared by hand picking twelve late L4 animals onto single plates. Unless noted, all animals carry the *fer*-*15(b26)* mutation, which causes a defect in spermatogenesis at 25°C that renders the animals self-sterile, facilitating the culture of age-synchronous animals. All live animals in a cohort were imaged at the same age. Animals were mounted for microscopy in a 4 μl drop of M9 buffer on a thin pad of 2% agarose containing 2 mM NaN_3_ and observed with a 60x and 100x Plan-Apochomat objectives (NA = 1.4) on a Nikon Eclipse 80i Microscope. Neurons were identified based on position and scored as missing when no fluorescence was observed within the cell body or its neurite.

### Lifespan assays

Lifespan assays were conducted as described^54^, without addition of 5-fluoro-2’-deoxyuridine (FUDR). After microscopy, animals were transferred singly onto plates. We used the L4 molt as age 0 for lifespan analysis. Assessment of viability and movement was performed every day until death, as described^30^.

### Computational power analysis

Simulations were performed in Matlab (Mathworks). We represented the correlation structure of lifespan and cephalic neuron-span using copulas. We generated random copulas with specific Spearman's rank correlations using the *copularnd* function. The marginal distribution of lifespans was the experimentally determined distribution of wild type or *age*-*1* mutants (3,397 and 542 animals, respectively). The marginal distribution of neuron-spans was not known, but the number of animals with a present or absent cephalic neuron on day 7 of adulthood was the experimentally determined. Using those values, we partitioned the simulated observations into “cell present” and “cell absent” groups, assigning the lowest neuron-span ranks to the latter. We performed this procedure 5,000 times for 75 Spearman's rank correlations values, which were more densely interspaced as the correlation magnitude approached zero. In each iteration, we calculated the mean lifespan of each group, and compared their survival curves using the Peto-Prentice Wilcoxon test. This enabled us to construct confidence intervals for mean lifespans, and to determine the fraction of tests that exhibited a significant difference at *P* < 0.05 (*i*.*e*. the statistical power of our experiment), as a function of the Spearman's rank correlation. Matlab scripts are available upon request.

### Statistical analysis

We used JMP 10 (SAS) software to carry out statistical analysis, to determine means and percentiles, and to perform ordinal logistic regressions. We determined whether the fractions of cells missing in two or more than two groups were equal using Fisher’s Exact test and Pearson’s chi-square test, respectively. We determined whether the survival functions of different groups were equal using the Wilcoxon test. *P*-values were calculated for control and experimental animals examined at the same time. We used the Bonferroni correction to adjust *P*-values when performing multiple comparisons.

## ACKNOWLEDGEMENTS

We thank Guy Caldwell for providing the *baInl1* strain; Jennifer Whangbo, Rebecca Ward and members of the Fontana lab for comments on the manuscript and stimulating discussions. Some of the *C. elegans* strains used in this work were provided by the *Caenorhabditis* Genetics Center, which is funded by US National Institutes of Health (NIH) Office of Research Infrastructure Programs (P40 OD010440). This work was funded by the NIH through grant R01 AG034994.

## AUTHOR CONTRIBUTIONS

J.A. Conducted experiments, simulations and statistical analysis. J.A. and W.F. conceived the project, interpreted data, and wrote the manuscript.

